# SIMPLE ASSAY FOR ADSORPTION OF PROTEINS TO THE AIR-WATER INTERFACE

**DOI:** 10.1101/2021.08.16.456550

**Authors:** Bong-Gyoon Han, Robert M. Glaeser

## Abstract

A rapid assay is described, based upon the Marangoni effect, which detects the formation of a denatured-protein film at the air-water interface (AWI) of aqueous samples. This assay requires no more than a 20 μL aliquot of sample, at a protein concentration of no more than1 mg/ml, and it can be performed with any buffer that is used to prepare grids for electron cryo-microscopy (cryo-EM). In addition, this assay provides an easy way to estimate the rate at which a given protein forms such a film at the AWI. Use of this assay is suggested as a way to pre-screen the effect of various additives and chemical modifications that one might use to optimize the preparation of grids, although the final proof of optimization still requires further screening of grids in the electron microscope. In those cases when the assay establishes that a given protein does form a sacrificial, denatured-protein monolayer, it is suggested that subsequent optimization strategies might focus on discovering how to improve the adsorption of native proteins onto that monolayer, rather than to prevent its formation. A second alternative might be to bind such proteins to the surface of rationally designed affinity grids, in order to prevent their diffusion to, and unwanted interaction with, the AWI.

## INTRODUCTION

With the emergence of single-particle electron cryo-microscopy (cryo-EM) as a popular technology for determining the structures of biological macromolecules (1), the ease with which proteins adsorb to the air-water interface (AWI) has become an issue of major concern (2). As background, cryo-EM uses specimens whose ideal thickness is comparable to the size of the macromolecular particles themselves, and in any case not thicker than about 100 nm. As a result, the time required for particles to diffuse to the AWI – after formation of a suitably thin specimen – is on the scale of 1 ms or less (3, 4). As a consequence, it is not possible for freely diffusing particles to avoid colliding many times with the AWI prior to vitrification.

Unfortunately, there is a high risk of denaturation upon contact, and thus touching the AWI is a very dangerous thing for proteins to do. Nevertheless, as many as 10% to 25% of samples can be prepared for cryo-EM with little difficulty, and preparation of many other samples, which may initially prove to be challenging, eventually succeeds after extensive, particle-specific optimization of the buffer and other conditions (5–7). One way to reconcile the frequent success with the obvious hazard of grid preparation might be to hypothesize that a sacrificial, denatured-protein skin almost always forms at, and passivates, the AWI, to which native proteins can be bound – at least under optimized conditions – in a structure-friendly way (2).

The literature provides surprisingly little guidance as to whether a newly investigated protein can be expected to denature upon diffusing to and touching the AWI. Although the study of protein adsorption to the AWI is quite mature, the number of test specimens that have historically been characterized is, in fact, rather limited relative to the number and variety of proteins now being studied by cryo-EM. Furthermore, the characterization of adsorption and denaturation of proteins at the AWI has historically involved a lengthy study in its own right, rather than employing a simple assay that could be used as part of a larger study. As a result, there is a need to develop fast and easy methods that might be used to assist optimization of the conditions used to prepare a new type of sample for cryo-EM.

Here we introduce an assay that uses the Marangoni effect (see section 20.4 of (8) for background) to detect the modification of the AWI that occurs when proteins are adsorbed to and denatured at the interface. Each test can be done in 10 minutes or less and requires only 10 μL or less of sample. Using this assay, we have observed that most proteins quickly modify the AWI, many at concentrations as low as 0.15 to 0.25 mg/mL. Two proteins, however, one of which has already been shown by cryo-EM to bind to the AWI with a strongly preferred orientation, produced relatively little effect, even at concentrations as high as 1 mg/mL.

We propose that this assay can, in the first instance, reduce or even remove uncertainly about whether a denatured-protein monolayer forms at the AWI, under the same conditions that are used when preparing grids for cryo-EM. This assay also provides an easy way to estimate the speed at which such denatured-protein monolayers are formed. Going beyond that, the assay provides a useful way to screen for additives or chemical treatments that might improve the preparation of EM samples by preventing adsorption of proteins to the AWI. The final proof of optimization will nevertheless remain with subsequently screening grids in the electron microscope.

## MATERIALS AND METHODS

### Reagents

A total of 13 soluble-protein samples were used as test specimens during the development and evaluation of the assay described here. The names of 11 of the soluble proteins, and the buffers in which they were tested, are given in **Table. 1**. Ferritin, carbonic anhydrase, lysozyme, cytochrome C, catalase, bovine serum albumin, and avidin were purchased from Sigma Aldrich, while streptavidin was purchased from New England Biolabs. Three additional soluble-protein samples were gifts of colleagues at Berkeley, as follows: rubisco, the lab of Prof. David Savage; FIP200:13, the lab of Prof. James Hurley; depolymerized tubulin, the lab of Prof. Eva Nogales.

Two highly polymerized soluble proteins, taxol-stabilized microtubules and Tobacco Mosaic Virus (TMV) respectively, were also tested with the assay. The taxol-stabilized microtubules, suspended in 80 mM PIPES, pH 6.8, with 1 mM EGTA, 1 mM MgCl2, 1 mM DTT, and 160 μM taxol, were again a gift of the lab of Prof. Eva Nogales, and the TMV sample was one that had been previously isolated by ourselves and stored as frozen aliquots.

In addition, the membrane protein ATG9 was tested, a gift of the lab of Prof. James Hurley. The buffer used for this sample was 50 mM HEPES, pH 7.5, with 150 mM NaCl, 0.02 dodecylmaltoside detergent (DDM) and 1 mM TCP.

Nonafluorobutyl methyl ether (NFBME) was purchased from Sigma Aldrich. NFBME is a non-flammable, low toxicity, volatile surfactant, see Appendix 5 of (9). We recommend it over other volatile surfactants that produce a desirable Marangoni effect, such as ethanol or chloroform, because of its extremely low solubility in water.

### Methods

The assay is performed by first applying 10 μL of an aqueous sample onto a hydrophilic surface in order to form a puddle that is ~1 cm in diameter. We use an ~1-inch square piece of freshly cleaved mica as the hydrophilic substrate, which produces a clean surface with minimal effort. A Hamilton syringe is then used to deliver a small drop, about 0.5 μL to 1.0 μL, of NFBME onto the middle of the puddle. The Marangoni effect, if any, already becomes apparent as the tip of the syringe approaches the thin puddle, but it becomes even easier to observe and record when the drop is deposited directly on top of the puddle of aqueous sample.

The assay is based upon the fact that bulk liquids flow in the direction of a gradient of surface tension. It is further based upon the supposition that a pre-existing monolayer of a particular surfactant that is contained with a sample, and which assembles at the air-water interface soon after spreading a thin film of sample, might effectively serve as a physical “cover slip” that seals the AWI and thus blocks the formation of a gradient in surface tension. Alternatively, the Marangoni effect might be suppressed if the viscosity of a pre-existing monolayer of surfactant is high enough.

The assay used here is based on observing the concentration of solute (e.g. protein) at which the Marangoni effect is completely abolished within a short, defined interval between first spreading the sample and subsequently depositing a drop of NFBME. We refer to this value of solute concentration as the “critical sealing concentration”, CSC, in analogy to the terminology used to refer to the concentration values at which various surfactants first assemble into micelles, i.e. the critical micelle concentration, or CMC. Unlike the formation of micelles, however, the assembly of a surfactant monolayer at the AWI is not a homogeneous process, and, for proteins, it is not expected to be reversible. As a result, the value of CSC must specify the time interval during which the monolayer is allowed to form. In the current work, the time interval is chosen – for practical convenience – to be ~10 s.

The setup that we have used for this assay is shown in **Figure 1**. The hydrophilic substrate (cleaved mica in our case) is placed on a dark background, as is shown in the photograph in panel A, but not in the cartoon in panel B, and the sample is viewed with light that is reflected at a shallow angle. Conditions of illuminating and viewing samples are adjusted empirically, in order to optimize the contrast produced as the shape of the puddle changes. Once the viewing angle and the illumination angle have been optimized, no further adjustment is needed, even during use on subsequent days. The specimen should be contained in a chamber that can be kept at high humidity, if it is desired to allow longer periods of time for a freshly prepared AWI to become modified. In our case the lid of the chamber is briefly removed to apply droplets of DFBME to the puddle of sample on the hydrophillic substrate and to observe the response.

**Figure 1.**
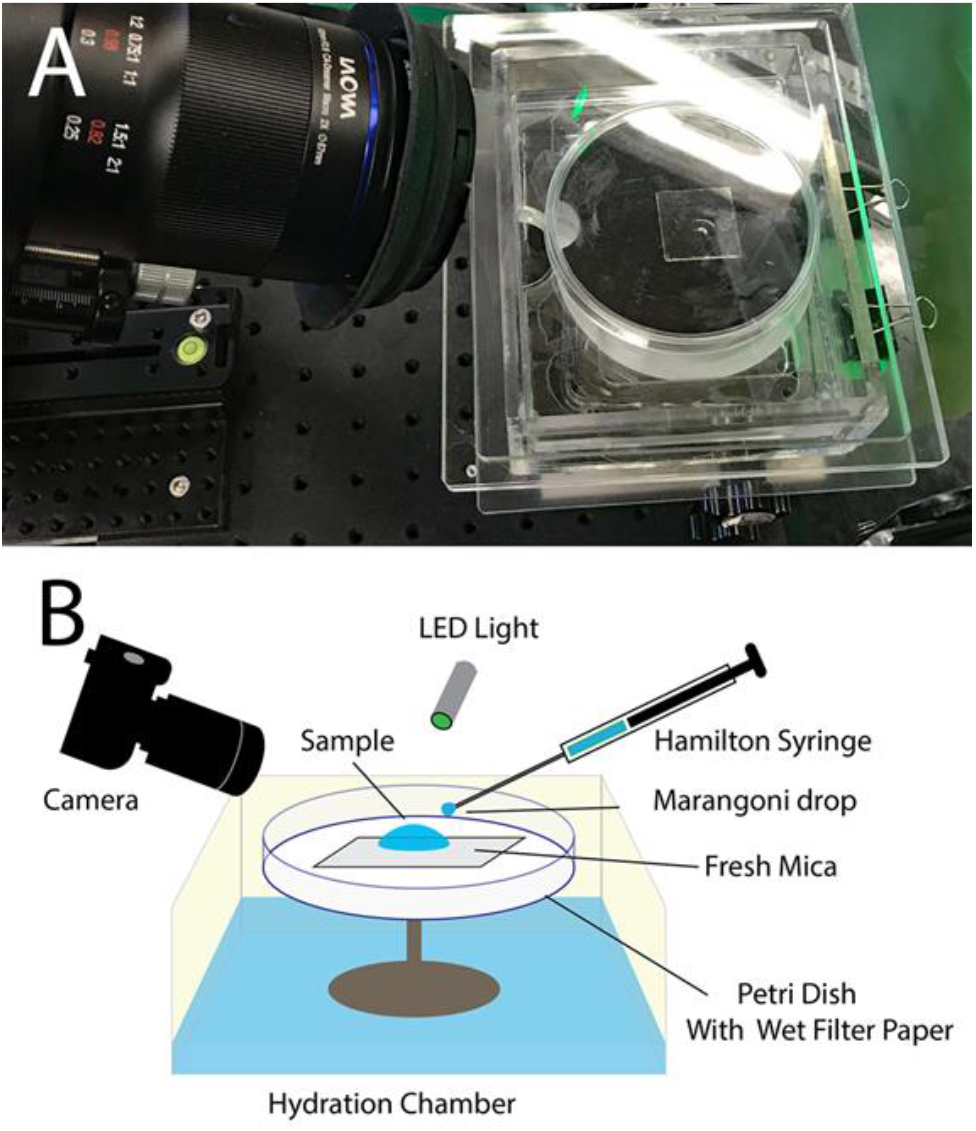
Setup to perform the assay. A hydrophilic substrate (cleaved mica in our case) is placed onto something that provides a dark background, as is shown in the photograph in **panel A**, but not in the cartoon in **panel B**, and the sample is viewed with light that is reflected at a shallow angle. Conditions of illuminating and viewing samples are adjusted empirically, in order to optimize the image contrast produced by the reflected light as the shape of the puddle changes, after which no further adjustment in the viewing and illumination angles is needed. The specimen is contained within a chamber that can be kept at high humidity, if it is desired to allow longer periods of time for a freshly prepared AWI to become modified, but the lid of the chamber is removed whenever DFBME droplets are applied to the puddle of sample on the hydrophilic substrate.

## RESULTS

When a small drop of nonafluorobutyl methyl ether (NFBME) is spotted onto a puddle of buffer, itself free of surfactant, this immediately causes the buffer to pull away from the point where the NFBME was applied. However, as the example in **Figure 2 F** illustrates, the buffer does not completely dewet the hydrophilic mica substrate, but, instead, a sheet of liquid remains that now is thin enough to exhibit interference fringes. The stability against dewetting suggests that NFBME, adsorbed at the AWI, generates a disjoining pressure that opposes thinning beyond a certain point. As further evidence that a film, or sheet, of liquid remains in the thinned area, the displaced puddle of buffer immediately returns to its initial shape and thickness as soon as the remaining droplet of NFBME evaporates (data not shown).

**Figure 2.**
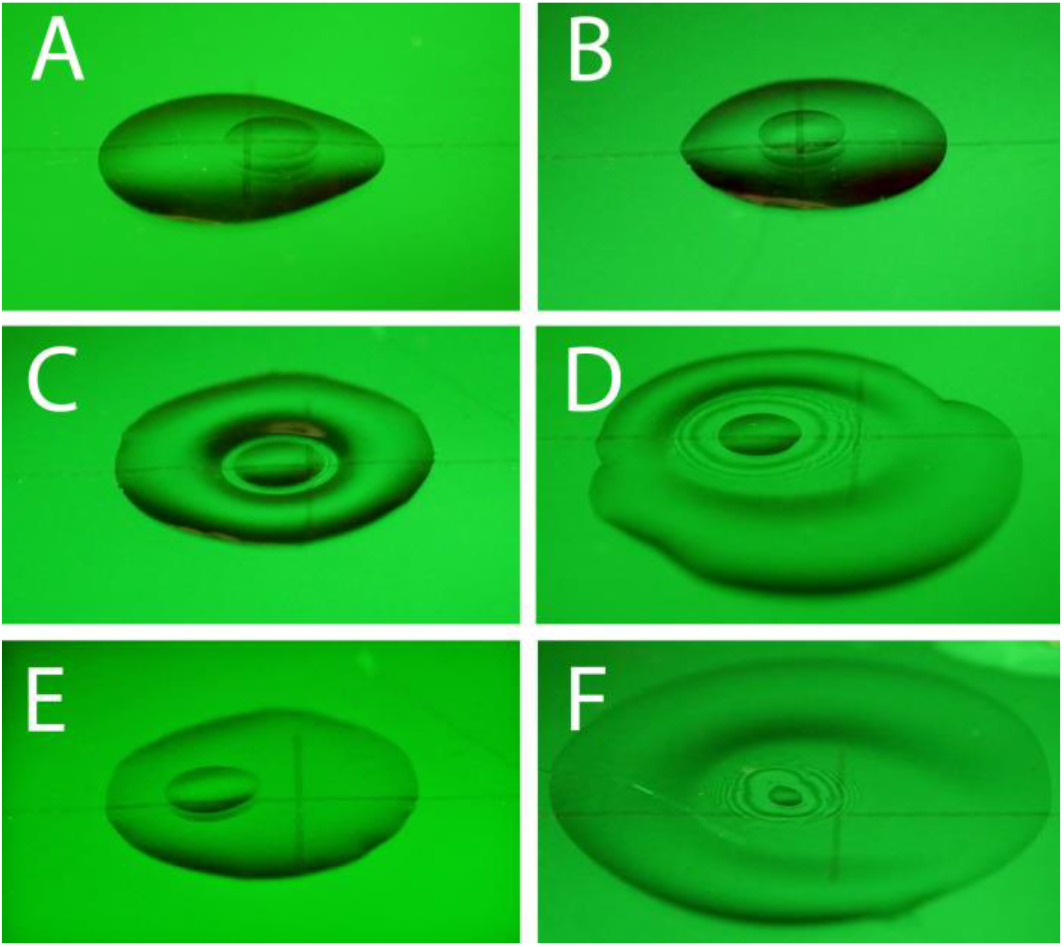
Responses of 10 μL aliqouts of ferritin samples at decreasing protein concentrations. **Panels A-D** show the responses seen within 10 s of applying the aliquots to the surface of freshly cleaved mica. The protein concentrations were (A) 0.45 mg/ml, (B) 0.15 mg/ml, (C) 0.05 mg/ml, and (D) 0.017 mg/ml, respectively. **Panel E** shows the response of the aliquot of 0.05 mg/mL ferritin solution after incubating for 10 minutes, and **panel F** shows the response of the buffer-only control.

The initial retraction of the body of the puddle of buffer, away from the small drop of NFBME, is an example of the Marangoni effect. In brief, the Marangoni effect represents the bulk movement of a liquid in response to there being a gradient in the surface tension of the liquid. The “tears of wine” phenomenon, the mechanism of which is explained in section 20.4 of (8), is another example of the Marangoni effect, and it is one that is familiar to many.

While the behavior of sufficiently dilute protein solutions is similar to that of buffer alone, at some point the Marangoni effect is no longer observed as the protein concentration is increased. Using ferritin as an example, **panels A – D in Figure 2** illustrate the concentration-dependence of the response that is observed within ~10 s of first applying 10 μL aliquots of sample onto freshly-cleaved mica. In this example, successive samples were prepared by 3-fold serial dilution of a concentrated stock. As is shown in **Figure 2**, the Marangoni effect is not evident at a ferritin concentration of 0.15 mg/mL, and it is already modified at a ferritin concentration of 0.05 mg/mL.

We hypothesize that the Marangoni effect is abolished at a ferritin concentration of 0.15 mg/mL because, at that concentration, a monolayer of denatured protein quickly forms at the AWI in less than 10 s. We further hypothesize that this monolayer of denatured protein effectively “seals” the top of the puddle of protein solution, as if a coverslip placed on top of the puddle, and thereby prevents the formation of a gradient in surface tension. Alternatively, we hypothesize that the monolayer may be so viscous, or even mechanically rigid, that no flow occurs. Either way, as already stated in the Methods section, we operationally define a “Critical Sealing Concentration (CSC)” as being the protein concentration at which the Marangoni effect is abolished within ~10 s of forming a fresh air-water interface.

Not surprisingly, we find that the values of CSC vary quite considerably for different types of proteins. **Figure 3** shows a second example, this time for serial 2-fold dilutions of a catalase sample, and **Table 1**. presents the CSC values for all protein samples included in this study. These values range from 0.15 mg/mL for ferritin, as is mentioned above, to 1 mg/mL for streptavidin and avidin. Two proteins, lysozyme and cytochrome C respectively, are too small to be imaged by cryo-EM at this time, but they are included here because they were frequently used as test specimens in early studies of protein denaturation at the AWI. As it happens, they both have CSC values, as measured by our assay, that are in the middle of the range covered by our complete set of protein samples.

**Figure 3.**
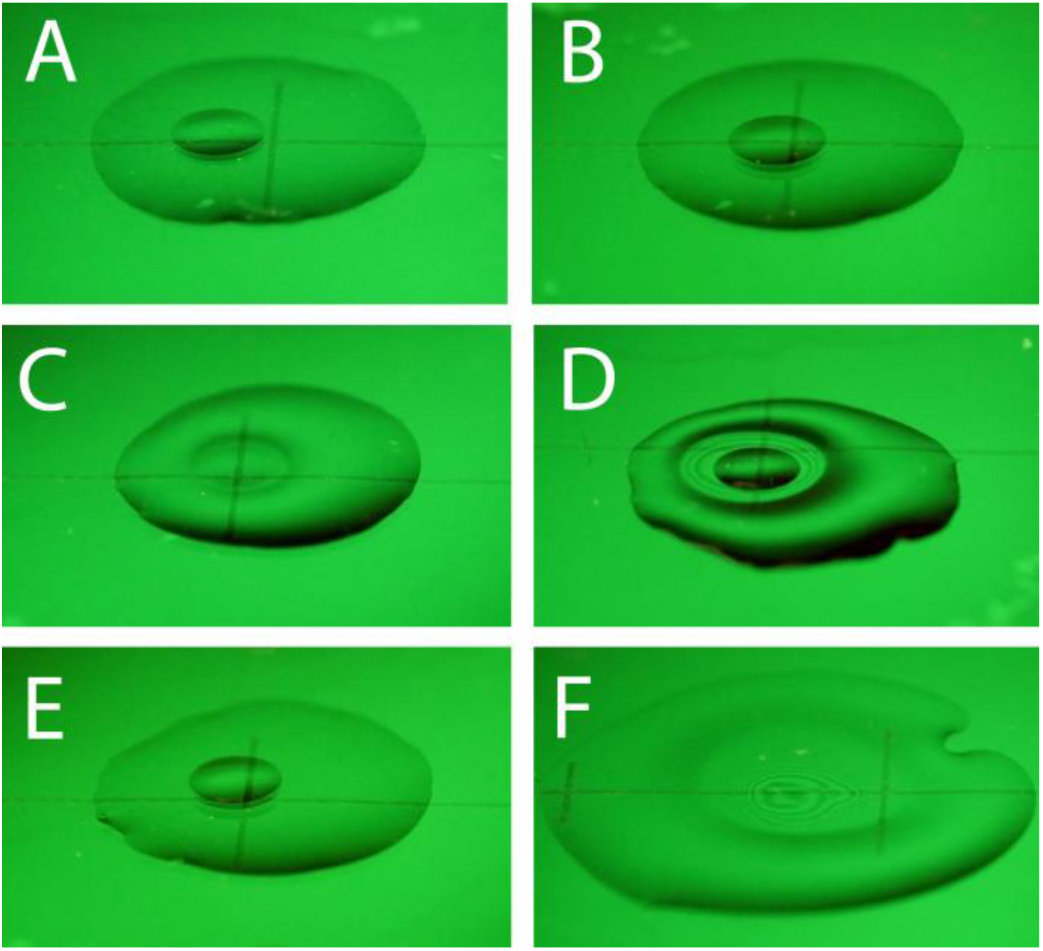
Responses of 10 μL aliqouts of catalase samples at decreasing protein concentrations. **Panels A-D** show the responses seen within 10 s of applying the aliquots to the surface of freshly cleaved mica. The protein concentrations were (A) 0.5 mg/ml, (B) 0.25 mg/ml, (C) 0.125 mg/ml, and (D) 0.0625 mg/ml, respectively. **Panel E** shows the response of the aliquot of 0.125 mg/mL catalase solution after incubating for 10 minutes, and **panel F** shows the response of the buffer-only control.

**Table 1.** Soluble proteins and buffers used to develop the assay described here, and values of critical sealing concentration (CSC) measured for each.

| **PROTEIN NAME** | **SAMPLE BUFFER** | **CSC (mg/mL)** |
| --- | --- | --- |
| **Ferritin** | **Buffer 1** | **0.15** |
| **Rubisco** | **Buffer 2** | **0.16** |
| **FIP200:13** | **Buffer 3** | **0.23** |
| **Tubulin (depolymerized)** | **Buffer 4** | **0.23** |
| **Carbonic anhydrase** | **Buffer 1** | **0.23** |
| **Catalase** | **Buffer 1** | **0.25** |
| **Lysozyme** | **Buffer 1** | **0.4** |
| **Cytochrome C** | **Buffer 1** | **0.5** |
| **Bovine serum albumin** | **Buffer 1** | **0.5** |
| **Streptavidin** | **Buffer 5** | **1.0** |
| **Avidin** | **Buffer 1** | **1.0** |

As might be expected, protein concentrations that initially exhibit only a reduced Marangoni effect, at the earliest time point, may progress to the point of eliminating it completely within ~10 minute. As examples, **Figures 2 C, E and 3 C, E** show the behaviors of ferritin and catalase solutions, respectively, at the initial time of ~10 s and then after waiting 10 min before applying a small drop of NFBME. As is addressed further in the Discussion, the formation of a denatured-protein layer at the AWI is expected to be a diffusion-limited process, but there may be steps, in addition to diffusion to the AWI, that limit the rate at which adsorbed proteins are converted into the molecularly thin equivalent of a coverslip.

While it is well known that detergents and other surfactants lower the surface tension of water, we were uncertain whether they might also form monolayers that act as a kind of coverslip, capable – in their own right – of interfering with the Marangoni effect. We find that this is, in fact, the case for NP40 and dodecyl maltoside (DDM) detergents, but not for CHAPSO. In the case of NP40 and DDM, both have CSC values of ~0.015 mg/mL that are well below their respective CMC values. In fact, their CSC values correspond to solution concentrations that contain just the amount of material needed, if almost fully partitioned to the AWI interface of the puddle, to form a ~1 nm-thick monolayer.

When a given type of detergent does block the Marangoni effect, it is logical to assume that the same detergent will also interfere with the use of the assay for membrane proteins. To test this expectation, we used the membrane protein ATG9 solubilized in DDM, a gift from the Hurley lab at Berkeley. While, as expected, we did not see a Marangoni effect, we were surprised to see – instead – that the ability of NFBMF to wet the surface of the puddle was increased considerably by the combination of detergent and protein. Importantly, the wettability of the same buffer is not increased when the AWI is sealed by detergent alone, as is seen in **Figure 4 F**, nor is the surface of a soluble-protein solution wetted by NFBME, as shown in **Figure 2 A and Figure 3 A**.

**Figure 4.**
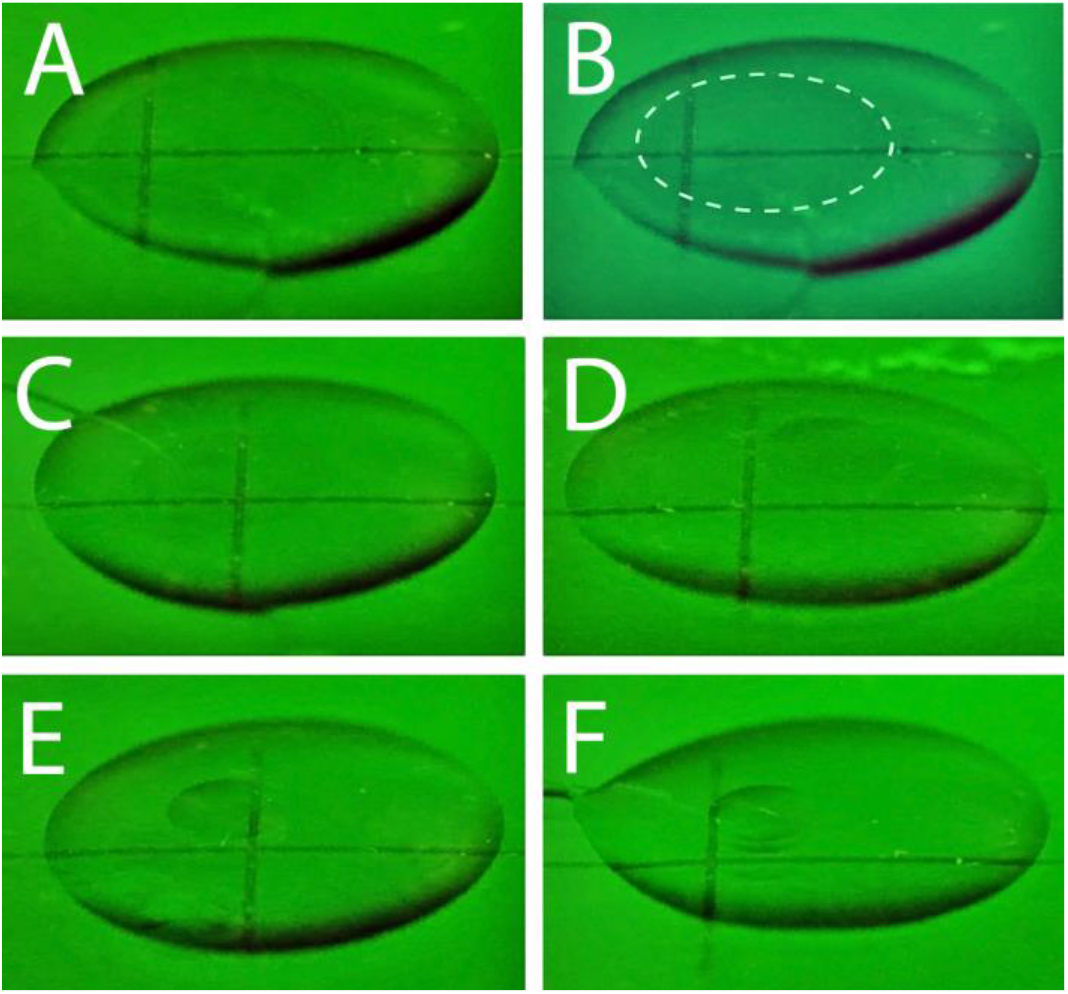
Responses of 10 μL aliquots of samples of the detergent-solubilized membrane protein, ATG9, at decreasing protein concentrations. **Panels A-E** show the responses seen within 10 s of applying the aliquots to the surface of freshly cleaved mica. The protein concentrations were (A) 0.85 mg/ml, (B) the same image as panel A, but with a dotted circle to indicate the area covered by DFBME, (C) 0.42 mg/ml, (D) 0.21 mg/ml, and (E) 0.11 mg/ml, respectively. **Panel F** shows the response of the detergent-containing, buffer-only control.

As **Figure 4** shows, the wettability of detergent-solubilized ATG9 increases with protein concentration, holding the DDM concentration constant. This new effect again suggests that protein must be inserted into the detergent monolayer, possibly displacing some of the bound detergent. While it is beyond the scope of the present study to investigate how general this new effect is for different detergents and for different types of protein, it will be of interest to determine whether the assay can be used for additional detergent-solubilized membrane proteins.

We also wanted to know whether linear protein polymers, such as TMV virus particles or taxol-stabilized microtubules, might be more resistant to the formation of a denatured-protein monolayer at the AWI. As is shown in **Figure 5**, this does seem to be true; although both polymers reduced the Marangoni effect within 10 s, at concentration values for which most other proteins seal the surface, they did not abolish it completely. Polymerized tubulin, at a protein concentration of 1 mg/mL, even retained a small Marangoni response after waiting for 10 minutes, as shown in **Figure 5 D**.

**Figure 5.**
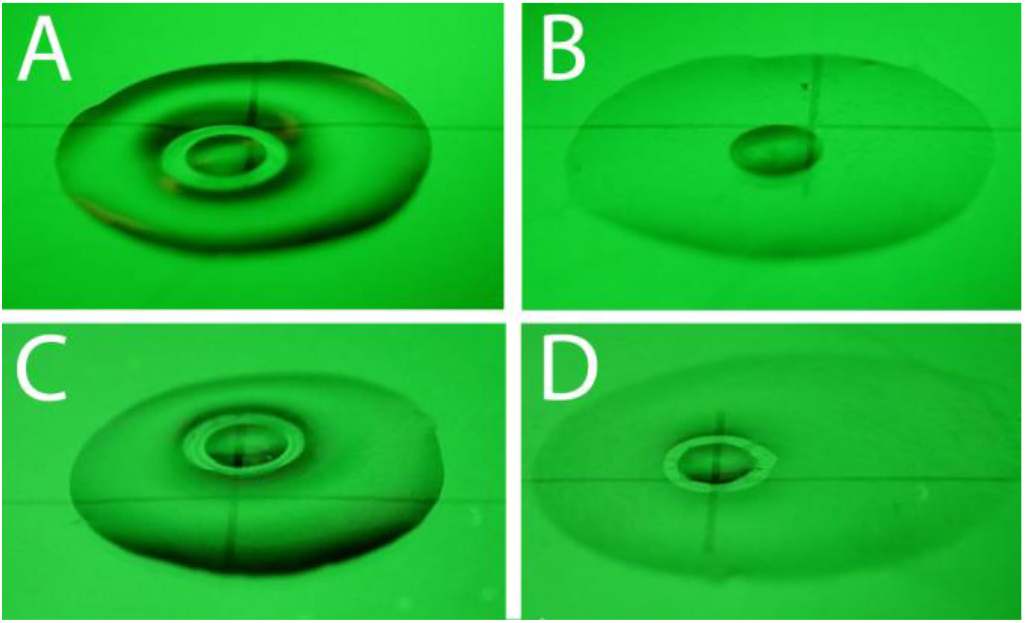
Responses of 10 μL aliquots of samples of TMV and taxol-stabilized microtubules, respectively. **Panels A and B** show samples of TMV at a concentration of 0.56 mg/ml, observed within 10 sec (A) and after 10 min (B). **Panels C and D** show samples of taxol-stabilized microtubules at a concentration of 1 mg/ml, observed within 10 sec (A) and after 10 min (B).

## DISCUSSION

The assay introduced here provides a fast and inexpensive way (A) to determine whether a specimen of interest makes a denatured-protein film at the air-water interface (AWI) and, if it does, (B) to estimate how rapidly such a film is formed. The assay is based upon the fact, demonstrated here, that local application of a water-insoluble, volatile surfactant causes a puddle of buffer to retract from the point of application, whereas moderately high concentrations of soluble proteins suppress or even abolish this effect. Retraction of buffer, which is an example of Marangoni flow, occurs because the volatile surfactant produces a gradient in surface tension. Our proposed explanation of suppression of the Marangoni effect is that a monolayer (or more) of adsorbed protein either blocks formation of such a gradient in surface tension, or, alternatively, it might form a stiff “coverslip” that prevents Marangoni flow. Either way, it is reasonable to expect that proteins that form such monolayers will be damaged, if not largely unfolded, soon after they adsorb to the AWI.

While most proteins are, in fact, likely to adsorb to the AWI, as was recently demonstrated by electron tomography (10), we find that they differ considerably in terms of the ease with which they suppresses the Marangoni effect. As is summarized in **Table 1**, ferritin and rubisco, at one extreme, block the Marangoni effect within 10 s at concentrations of only ~0.15 mg/mL. Streptavidin and avidin, at the other extreme, reduce but do not fully block the Marangoni effect at ~7 times higher protein concentration (1 mg/mL). Depolymerized tubulin blocks the Marangoni effect quite easily, but polymerized tubulin is much less effective in doing so. Interestingly, proteins that have been used extensively in studies of protein denaturation at the AWI (lysozyme, cytochrome C, and bovine serum albumin) are among those that require higher concentrations to rapidly block the Marangoni effect.

Diffusion to the AWI is the first, but not necessarily the only factor that limits the rate at which a complete film of adsorbed protein is formed. It is not surprising, therefore, that how quickly the Marangoni effect is blocked in this assay depends upon the protein concentration. Other factors that might affect the rate include (A) the value of the sticking coefficient, i.e. the fraction of collisions that, on average, actually result in adsorption of proteins; (B) how quickly and how extensively a given type of protein unfolds after it binds to the surface; and (C) how fully the surface must be covered before the Marangoni effect is finally blocked. It is not surprising, therefore, that the values of concentration for which Marangoni flow is already blocked within about 10 s, i.e. the different CSC values listed in **Table 1**, vary by almost a factor of 10.

In most but not all cases, the CSC value is much lower than the protein concentrations that are commonly used to prepare grids for cryo-EM, i.e. ~1.0 mg/mL. While formation of a denatured-protein monolayer at those concentrations is expected to happen too fast to be easily observed by this assay, one can nevertheless use the measured value of CSC to estimate, if only roughly, how quickly such a monolayer might form at a higher concentration.

A first such estimate might assume that the process is diffusion limited, in which case the time that it takes to deliver the needed amount of protein to the AWI will scale as the square of the distance from which the protein must be recruited, and thus as the square of the concentration. In addition to this, the time needed to seal the AWI will scale inversely with the sticking coefficient, and, as mentioned above, the sticking coefficient may vary from one protein to another.

Since the “CSC of the sample” is defined to be the concentration for which Marangoni flow is blocked within ~10 s, one can use the simple equation

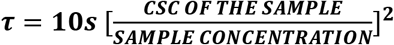

to calculate an upper bound for the time, **τ**, that sealing the AWI might take at the sample concentration one actually uses to prepare grids. For example, solutions of ferritin or rubisco, at a concentration of 1 mg/ml, are expected to be fully sealed within ~200 ms. Taking the calculation even further, it is expected that enough ferritin molecules would be adsorbed within 25 ms to modify the AWI to the same extent that is shown in **Figure 2C**. It must be realized, however, that using this equation is not appropriate if other factors are rate-limiting, such as unfolding or rearranging proteins at the AWI,.

One might initially think that properly folded proteins would actually avoid interaction with the AWI, because their surfaces, being largely hydrophilic, ought to resist dewetting. On the other hand, proteins generally have small hydrophobic patches on their surfaces, which may easily dewet upon collision with the AWI. As hypothesized previously (11), the resulting, initial-adsorption event may do little or no structural harm, other, perhaps, than causing preferential orientation. However, small portions of individual protein domains, referred to as “foldon units” (12), are thought to continuously fluctuate between folded and unfolded states. As a consequence, adsorption is expected to set the stage for a thermodynamically downhill cascade leading to very rapid, irreversible denaturation (11). We can thus imagine how the entire AWI would eventually become covered with a sacrificial, denatured-protein monolayer, to which additional protein molecules might subsequently adsorb while retaining their native structure,.

If a protein of interest does form a denatured-protein film at the AWI, as confirmed by using this assay, and if the this happens too fast to be outrun by rapid vitrification (7), there can still be a number of ways to optimize the preparation of grids for single-particle cryo-EM. Formation of a “coverslip” that passivates the AWI, as was envisioned by (13) to occur in the case of lipids, may even be a good thing. The possibility exists, for example, that denatured-protein coverslips are responsible for many if not all good results, provided that they are structure-friendly to the remaining particles in solution, and provided that they do not bind particles in preferred orientations. One option, therefore, is to modify the strengths of the interactions by which healthy proteins adsorb to a sacrificial monolayer, so as to suppress preferential orientation, further disassembly, or even formation of amorphous aggregates. For example, chemical modification of amino or carboxyl groups on the surfaces of proteins can be expected to modify the electrostatic interactions between well-folded proteins and sacrificial films. This could be a factor – along with crosslinking – that explains why reacting with glutaraldehyde or BS3 sometimes optimizes the preparation of grids. Similarly, detergents or other, small-molecule amphiphiles and surfactants may bind to hydrophobic patches on the surfaces of folded proteins, as well as modifying the chemical composition of any monolayers that passivate the AWI, and either effect may lead to optimization of grids prepared for cryo-EM.

Detergents are also expected to suppress the Marangoni effect, and thus they can interfere with using this assay for solubilized membrane proteins. Unexpectedly, however, we found that the wettability of the AWI interface by NFBME was increased for the sample of a detergent-solubilized membrane protein that was made available for doing a preliminary test. This effect opens the possibility that changes in wettability may extend the variety of specimens for which the assay could guide the successful preparation of grids for cryo-EM.

Finally, we point out that immobilization of particles onto biochemically functionalized EM grids offers an alternative way to prepare specimens for cryo-EM, and it is one that can be more rationally controlled – see Section 3 of (2). Streptavidin affinity grids have been used, for example, to obtain structures of particles for which other optimization methods had previously failed (14–16). While immobilization onto structure-friendly support films can prevent particles from making contact with the AWI, such support films do add a structural background that may make it more difficult to obtain high-resolution structures of smaller particles. Even so, functionalized graphene oxide grids have been used to obtain the structure of TRAP1 dimer, whose particle weight is only ~165 kDa (17), thus pointing to the possibility that added structural background may not be as limiting as has been supposed.

## ACKNOWLEDGEMENTS

This work has been supported in part through funds awarded by Genentech, and we thank Dr. Chris Arthur and others at Genentech for their interest in our work on protein adsorption to the AWI. We thank the following colleagues at the University of California for donating samples to use in the development and testing of the assay described here: samples of rubisco were provided by Luke Oltrogge in the laboratory of Prof. David Savage; samples of FIP200:13 and ATG9 were provided by Adam Yokum in the laboratory of Prof. James Hurley; and samples of taxol-stabilized microtubules and of depolymerized tubulin were provided by Julia Peukes in the laboratory of Prof. Eva Nogales.

